# Synthetic blood-based infrared molecular fingerprints: artificial cohorts for methodological research

**DOI:** 10.1101/2025.10.27.684739

**Authors:** Niklas Leopold-Kerschbaumer, Nico Feiler, Kosmas V. Kepesidis

## Abstract

Infrared molecular fingerprinting of human blood samples provides a powerful, minimally invasive approach for disease detection and health monitoring. However, ethical and legal constraints often limit the sharing of real patient data collected from clinical studies. In this work, we present a synthetic dataset of blood-based infrared molecular fingerprints, generated using multivariate Gaussian models fitted on real measurements from a large case-control study targeting various cancer types. The synthetic dataset retains the statistical and physical properties of real molecular fingerprints, enabling the development and validation of analytical methodologies without compromising patient privacy. We demonstrate that the provided artificial dataset can serve as a proxy for real data in methodological research, facilitating reproducibility and collaboration in biomedical spectroscopy. This approach offers a practical solution for overcoming ethical barriers in clinical data sharing in spectroscopic biomarker research.

## BACKGROUND & SUMMARY

Advancements in spectroscopic techniques have opened new frontiers in minimally-invasive disease diagnostics and personalized health monitoring ^1–4^. Among these, infrared molecular fingerprinting of blood-based samples has demonstrated significant potential for detecting and characterizing various pathological conditions, including cancer and metabolic disorders ^5–16^. However, the full utility of such datasets is often hindered by strict ethical and legal restrictions that prevent their open sharing, thereby limiting collaborative research and methodological development.

To address this challenge, we generate synthetic blood-based infrared molecular fingerprints that retain the essential statistical and physical properties of real patient data. Synthetic datasets are constructed using multivariate Gaussian models fit on real-world measurements from a cross-sectional study investigating various cancer types. By capturing and reproducing key spectral patterns in an artificial yet statistically valid manner, the synthetic cohorts provide a valuable resource for researchers aiming to develop and refine analytical techniques without direct access to sensitive patient data.

This paper presents the methodology used to generate the synthetic dataset, evaluates its fidelity to real-world spectral fingerprints, and discusses its potential applications in biomedical research. Importantly, the resulting synthetic cohorts are made publicly available through a dedicated open-access repository, enabling immediate and unrestricted use by the research community ^17^. This approach facilitates ethical data sharing, reproducibility, and innovation in the field of biomedical spectroscopy. Our findings support the broader adoption of synthetic datasets as a viable solution for overcoming ethical barriers while advancing the study of blood-based molecular diagnostics.

## METHODS

### Infrared fingerprinting of human blood plasma

Blood samples utilized in this work were collected in the framework of the multi-center *Lasers4Life* clinical study. This study was conducted in the Munich area and is registered (ID DRKS00013217) at the German Clinical Trials Register (DRKS). The study was reviewed and approved by “Ethikkommission bei der LMU München” (EK 20170820—Nr.:17-532), and was conducted according to Good Clinical Practice (ICH-GCP), the principles of the Declaration of Helsinki, and all applicable legislations and regulations. Informed consent was obtained from all participants prior to blood collection. Blood plasma samples of study participants were obtained following previously established sample handling procedures ^5^. Before measuring the samples using a commercially available FTIR device (MIRA-Analyzer, CLADE GmbH), the obtained blood samples were split into a training set and a test set. The measurements were carried out in a fully randomized manner over 19 weeks. After a 10-week gap, introduced to account for potential drifts in spectrometer performance and ensure robust testing, the test set was measured in randomized order over 2 weeks. The infrared molecular fingerprints collected span the spectral range of 930-3051 cm^-1^, capturing characteristic absorption bands for proteins, carbohydrates, and lipids. Once all measurements were obtained, outliers were detected using the Local Outlier Factor method from the scikit-learn library in Python 1.6.1, separately for the training and the test set. This ensured that erroneous or incomplete measurements were removed before analysis.

### Case-control designs

The health status of individuals in the study is categorized into either therapy-naïve patients with lung, prostate, bladder, or breast cancer or non-symptomatic reference individuals. Based on these classifications and available demographic information, case-control designs were constructed to ensure meaningful comparisons. To reduce potential confounding effects, statistical matching was performed using propensity scores, aligning cases with their closest control counterparts based on age and sex. The matching algorithm was implemented in R version 4.4.1. Following this procedure, the resulting matched cohorts comprise a total of 2,079 individuals, with 1,650 assigned to training sets and 429 to test sets. Detailed information on cohort demographics and sample sizes is provided in Table S1 in Supplementary Information A. Using these well-balanced cohorts, we fitted statistical models capable of generating new synthetic infrared spectra that capture the key statistical and spectral properties of the original datasets.

### Multivariate Gaussian modeling

Assuming that absorbance values at each wavenumber are approximately Gaussian distributed, synthetic infrared spectra can be generated by sampling from a multivariate Gaussian distribution (MVG) fitted to measured spectra. The MVG defines the probability density function

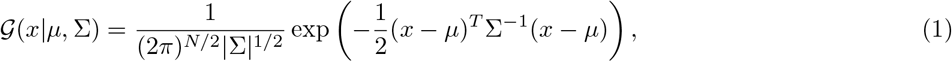

where *µ* ∈ ℝ^*N*^ represents the mean spectrum, and Σ ∈ ℝ ^*N ×N*^ is the covariance matrix capturing spectral correlations. Both the mean vectors and covariance matrices are provided for each dataset split described in Section . This modeling approach builds on prior work that applied MVG distributions to generate synthetic FTIR spectroscopy datasets for purposes such as sample size planning and method development ^18–22^. In addition to patient spectra, MVG models were also fit to quality control (QC) samples—constructed by pooling blood specimens—with 1,048 QC samples included in the training set and 131 in the test set.

## SYNTHETIC DATA RECORDS

Using our approach, we generated synthetic spectra for matched cohorts of healthy and diseased individuals across several cancer types: lung cancer (luca), breast cancer (brca), bladder cancer (blca), and prostate cancer (prca). For lung cancer, additional stratification was performed by disease stage, resulting in subcohorts for stages I through IV. Due to limited data for early-stage lung cancer, we also include a multivariate Gaussian (MVG) fit for a combined cohort representing stages I and II.

As previously described, each (sub-)cohort was divided into training and test sets prior to sample measurement. Within each set, samples were further grouped by disease status (healthy or diseased) and sex (male or female). A schematic overview of the full data simulation pipeline is shown in Figure 1A, while detailed data splits for each (sub-)cohort are presented in Figure 1B. Every data split highlighted in orange in Figure 1B corresponds to a fitted MVG model—characterized by a mean spectrum and covariance matrix—available in our GitHub repository ^17^.

**Figure 1.**
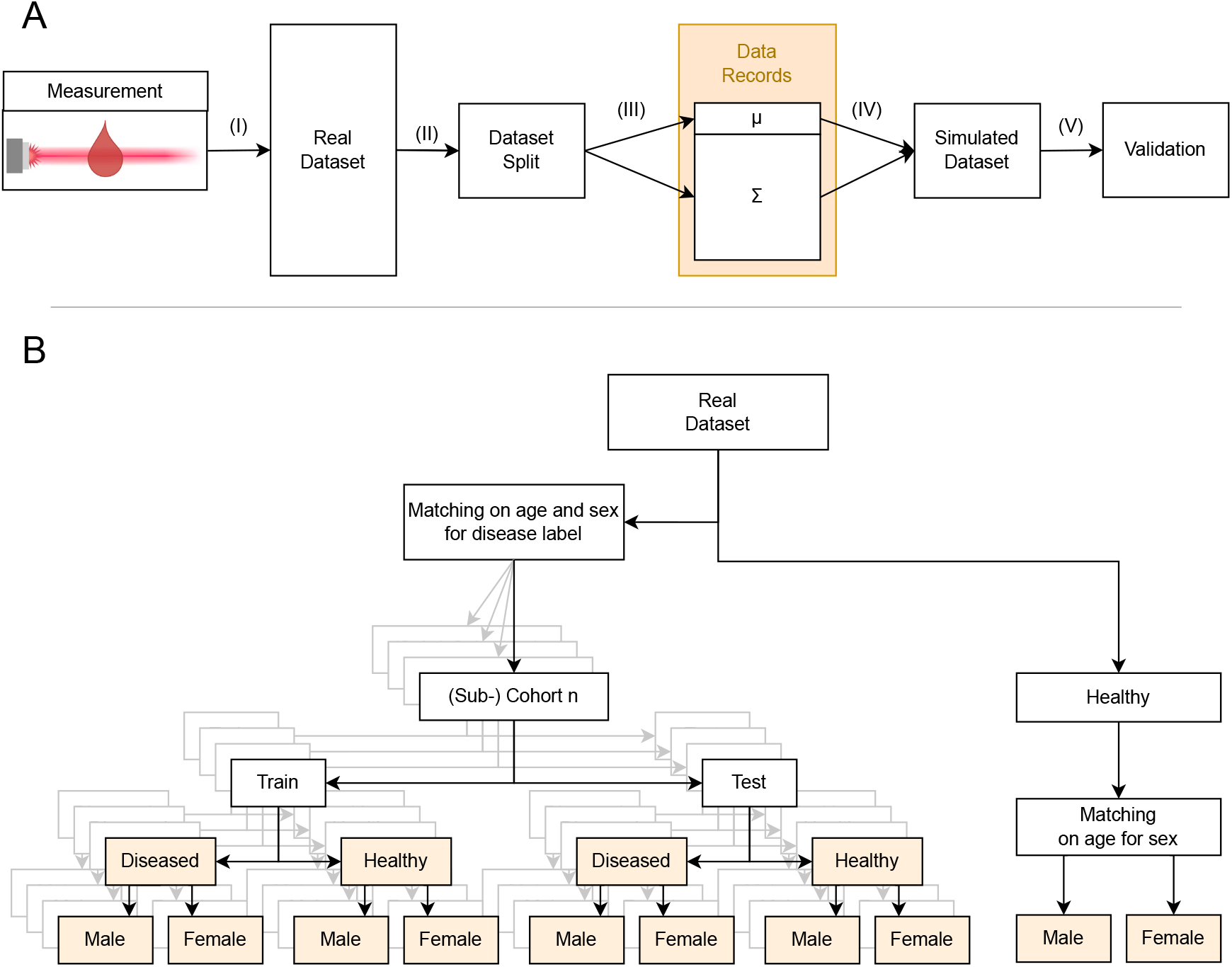
**A)** Diagram illustrating how the simulated datasets are obtained. (I) Samples of study participants are measured via FTIR, and measurements are stored in a dataset. (II) The real dataset is split into subsets based on disease status and/or sex according to Figure 1. (III) The mean and covariance matrix are calculated for each subset. (IV) Simulated spectra are sampled according to Equation 1. (V) The quality of simulated datasets is verified by comparing the ROC curves, differential fingerprints, and effect sizes to real data. **B)** Dataset splits visualized. Orange colored boxes indicate that the mean and covariance matrix are available for sampling new spectra using an MVG.

These MVG models, defined by Equation 1, enable users to sample synthetic spectra conditioned on sex (male or female), disease status (healthy or diseased), and for lung cancer, disease stage (I–IV). Additionally, we provide a reference cohort composed solely of healthy individuals, stratified by sex, from which MVG models for healthy male and female spectra were derived.

In the following section, we validate this simulation approach with use cases motivated by real-world applications such as disease prediction.

## TECHNICAL VALIDATION

To validate our simulation approach, we simulated cohorts of equal sizes to the real data and compared effect sizes (Cohen’s d) by analyzing both the full spectra and selected peak ratios, in line with prior work ^5^. We also evaluated the performance of a logistic regression classifier with L2 regularization (*C* = 10) on both datasets, using ROC curves for comparison. Results for the lung cancer cohort, stratified by disease status and sex (fourth row in Figure 1), are summarized in Figure 2.

**Figure 2.**
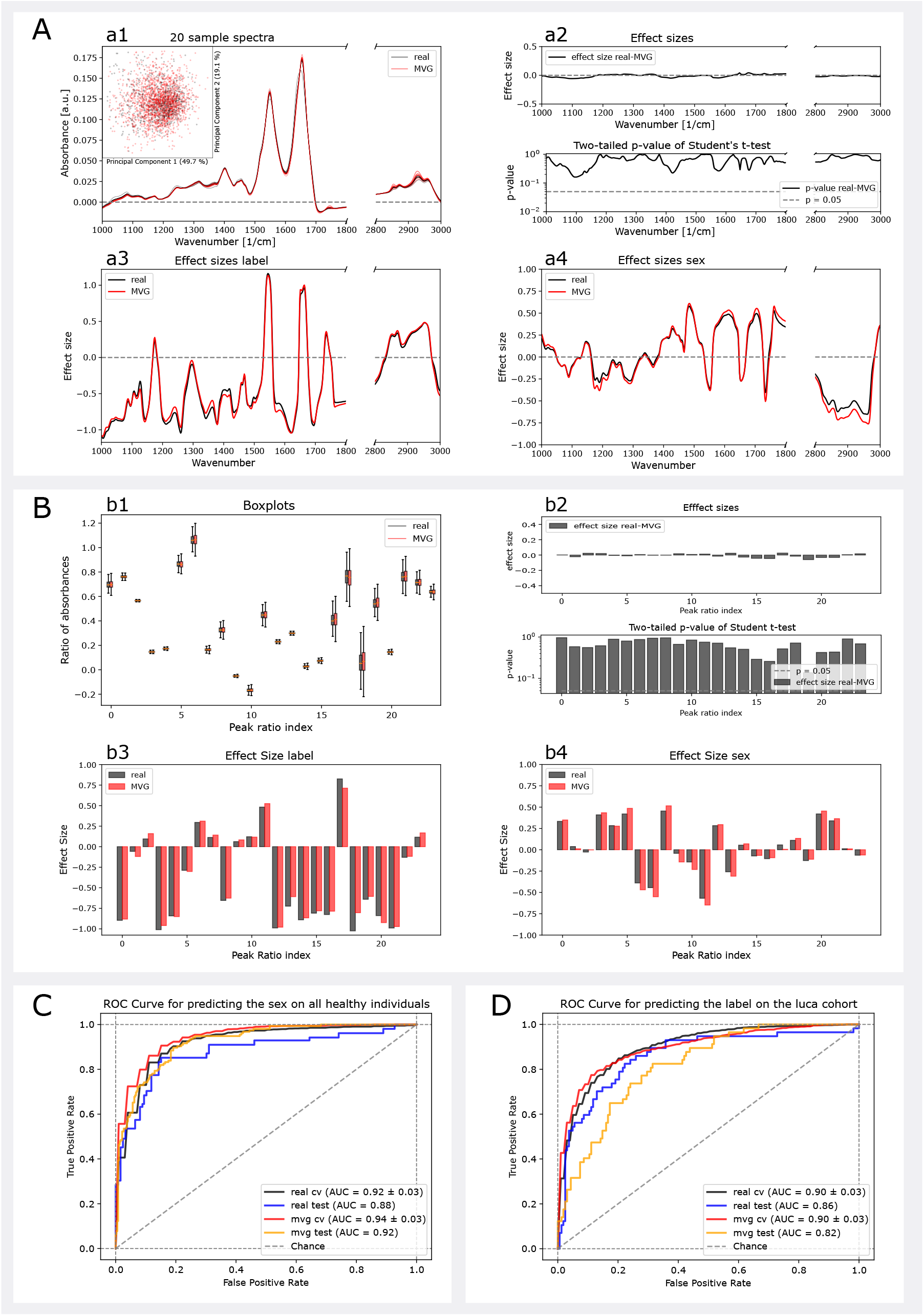
Comparison between real data and simulated data. **A)** Basic analysis on the spectra including 1. 10 spectra of real data and 10 spectra of simulated data with the outlier removed, PCA scatter plot of all spectra, 2. Effect size and t-test p-values between real and simulated data, 3. Effect size between the lung cancer label for real and simulated data, 4. Effect size between the sex for real and simulated data. **B)** Basic analysis on the peak ratios, including 1. Boxplot for each peak ratio for real and simulated data, 2. Effect size and t-test p-values between real and simulated data, 3. Effect size between the lung cancer label for real and simulated data, 4. Effect size between the sex for real and simulated data. **C)** ROC curve for predicting sex with the logistic regression classifier described above on all healthy individuals on the test set as well as within the training set for both real and simulated data. **D)** ROC curves for predicting the disease with the logistic regression classifier described above on the lung cancer cohort on the test set as well as within the training set for both real and simulated data.

Figure 2a1 shows 10 representative spectra alongside a PCA scatter plot, illustrating the visual similarity between real and simulated data. As seen in Figure 2a2, the effect size between real and simulated datasets is close to zero across all wavenumbers. Similarly, a test comparing the means per wavenumber shows that there is no significant difference between real and simulated data. Figures 2a3 and 2a4 demonstrate that effect sizes with respect to disease label and sex in the simulated data closely mirror those in the real data.

Similarly, Figure 2b1 shows that the distribution of peak ratios in the simulated data closely matches that of the real data. Figures 2b2–b4 provide a corresponding effect size and t-test analysis on these peak ratios, analogous to panel A. Which peak ratios were assigned to which index is shown in S2 in *Supplementary Information B*.

In terms of classification performance, the ROC curves in Figure 2C and Figure 2D show comparable results for sex prediction on all healthy individuals and disease prediction on the lung cancer cohort across real and simulated data, as well as between the training and independent test sets. The results also reveal a lower AUC for classification in the test set compared to the training set, indicating a domain shift. Notably, classifiers trained on multivariate Gaussian (MVG) simulations generally slightly outperform those trained on real data, while performance decreases on the independent test set. A complete summary of AUC scores across all dataset splits using the logistic regression classifier is provided in *Table S3* in *Supplementary Information C*.

All data underwent the following preprocessing steps, consistent with established methods ^6,12^:

### 1. Truncation I

Spectra were truncated to the range of 1000–3000 cm^-1^ to standardize measurements, excluding regions without peaks.

### 2. L2 Normalization

Each spectrum was normalized by its L2 norm:

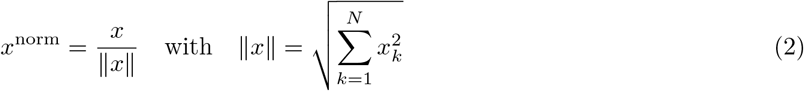

### 3. Truncation II

Absorbance values and wavenumbers between 1800–2800 cm^-1^ were removed. This “silent region” is dominated by water absorbance and contains no no molecular information.

For training logistic regression models, we applied standard scaling to the preprocessed data. The ROC curves on the training set were calculated using repeated cross-validation with 10 splits and 5 repetitions, while for evaluating the test set the entire training set was used. This setup, along with the preprocessing pipeline and classification model, follows established practices in blood-based FTIR spectroscopy tasks ^6^.

All technical validation experiments were conducted using Python 3.10.16 with NumPy 2.2.4 and scikit-learn 1.6.1.

## USAGE NOTES

### Limitations

While simulations based on multivariate Gaussian (MVG) fits provide highly interpretable approximations of data distributions, they come with important limitations. Most notably, MVG-based sampling does not support accurate modeling of individual-level variability. As a result, the generated spectra do not account for personal covariates such as age or BMI. Additionally, because the underlying dataset originates from a case-control study, it does not support longitudinal simulations or temporal analyses. Finally, MVG models assume linear relationships among features; any non-linear dependencies across wavenumbers present in real measurements may be lost in the fitted distributions.

### Prospects

If promising results are achieved using the simulated data, researchers are encouraged to contact the corresponding authors to explore validation opportunities on real datasets.

## DATA AVAILABILITY

All mean spectra and covariance matrices described previously are available in our GitHub repository ^17^.

## CODE AVAILABILITY

The GitHub repository ^17^ also includes an example Python script demonstrating how to use these models to generate synthetic spectra.

## AUTHOR CONTRIBUTIONS

**Niklas Leopold-Kerschbaumer:** Investigation; Formal analysis; Visualization; Methodology; Writing – original draft preparation; Writing – review & editing. **Nico Feiler:** Investigation; Formal analysis; Visualization; Methodology; Writing – original draft preparation; Writing – review & editing. **Kosmas V. Kepesidis:** Conceptualization; Supervision; Project administration; Methodology; Writing – original draft preparation; Writing – review & editing.

## ACKNOWLEDGMENTS

We thank Mihaela Žigman and all contributors involved in the design and execution of the *Lasers4Life* clinical study. We also express our gratitude to Ferenc Krausz for fostering the research environment that enabled the realization of this work.

## References

[1] Baker, M. J. et al. Using fourier transform ir spectroscopy to analyze biological materials. Nature protocols 9, 1771–1791 (2014).

[2] Butler, H. J. et al. Development of high-throughput atr-ftir technology for rapid triage of brain cancer. Nature communications 10, 4501 (2019).

[3] Voronina, L. et al. Molecular origin of blood-based infrared spectroscopic fingerprints. Angewandte Chemie 133, 17197–17206 (2021).

[4] Paraskevaidi, M. et al. Clinical applications of infrared and raman spectroscopy in the fields of cancer and infectious diseases. Applied Spectroscopy Reviews 56, 804–868 (2021).

[5] Huber, M. et al. Stability of person-specific blood-based infrared molecular fingerprints opens up prospects for health monitoring. Nature communications 12, 1511 (2021).

[6] Huber, M. et al. Infrared molecular fingerprinting of blood-based liquid biopsies for the detection of cancer. Elife 10, e68758 (2021).

[7] Martin, F. L. et al. Distinguishing cell types or populations based on the computational analysis of their infrared spectra. Nature protocols 5, 1748–1760 (2010).

[8] Ghimire, H. et al. Protein conformational changes in breast cancer sera using infrared spectroscopic analysis. Cancers 12, 1708 (2020).

[9] Ollesch, J. et al. An infrared spectroscopic blood test for non-small cell lung carcinoma and subtyping into pulmonary squamous cell carcinoma or adenocarcinoma. Biomedical Spectroscopy and Imaging 5, 129–144 (2016).

[10] Eissa, T. et al. Plasma infrared fingerprinting with machine learning enables single-measurement multi-phenotype health screening. Cell Reports Medicine 5 (2024).

[11] Kepesidis, K. V. et al. Assessing lung cancer progression and survival with infrared spectroscopy of blood serum. BMC medicine 23, 101 (2025).

[12] Kepesidis, K. V. et al. Breast-cancer detection using blood-based infrared molecular fingerprints. BMC cancer 21, 1–9 (2021).

[13] Backhaus, J. et al. Diagnosis of breast cancer with infrared spectroscopy from serum samples. Vibrational Spectroscopy 52, 173–177 (2010).

[14] Elmi, F., Movaghar, A. F., Elmi, M. M., Alinezhad, H. & Nikbakhsh, N. Application of ft-ir spectroscopy on breast cancer serum analysis. Spectrochimica Acta Part A: Molecular and Biomolecular Spectroscopy 187, 87–91 (2017).

[15] Ollesch, J. et al. It’s in your blood: spectral biomarker candidates for urinary bladder cancer from automated ftir spectroscopy. Journal of biophotonics 7, 210–221 (2014).

[16] Anderson, D., Anderson, R., Moug, S. & Baker, M. Liquid biopsy for cancer diagnosis using vibrational spectroscopy: systematic review. BJS open 4, 554–562 (2020).

[17] Leopold-Kerschbaumer, N. MVG-IMF. https://github.com/Attoworld-Data-Science/MVG-IMF (2025).

[18] Beleites, C., Neugebauer, U., Bocklitz, T., Krafft, C. & Popp, J. Sample size planning for classification models. Analytica chimica acta 760, 25–33 (2013).

[19] Eissa, T., Kepesidis, K. V., Zigman, M. & Huber, M. Limits and prospects of molecular fingerprinting for phenotyping biological systems revealed through in silico modeling. Analytical Chemistry 95, 6523–6532 (2023).

[20] Nemeth, F. B. et al. Bridging spectral gaps: Cross-device model generalization in blood-based infrared spectroscopy. Analytical Chemistry (2025).

[21] Wegner, C. et al. Toward informative representations of blood-based infrared spectra via unsupervised deep learning. Journal of Biophotonics e70011 (2025).

[22] Schroeder, J. et al. Information-optimal measurement: From fixed sampling protocols to adaptive spectroscopy. arXiv preprint arXiv:2505.14364 (2025).

