## Supporting Information for "Synthetic blood-based infrared molecular fingerprints: artificial cohorts for methodological research"

Additional information including (A) the overview of pipeline to get simulated datasets, (B) Table about the basic demographics of all (sub-)cohorts, (C) Table of all AUCs for real and simulated datasets for all (sub-)cohorts for disease label classification or sex classification (if applicable).

### Contents

**A Demographics of cohorts** **S3**

**B Peak ratio indices** **S4**

**C AUCs for all FTIR datasets** **S5**

### List of Tables

S1 Demographics across different diseases, stages, and train/test sets. . . . . . . . . . . . . . . . . . . . . . . S3
S2 List of Peak Ratios and assigned indices in Figure 2 . . . . . . . . . . . . . . . . . . . . . . . . . . . . . S4
S3 AUCs for predicting the disease label and the sex on real and on simulated data on all cohorts. . . . . S7

### A Demographics of cohorts

| Disease | Set | Size | Population (%) | | | | Age | | BMI | |
|---|---|---|---|---|---|---|---|---|---|---|
| | | | Healthy | | Diseased | | | | | |
| | | | Male | Female | Male | Female | Avg | Std | Avg | Std |
| BLCA | Test | 213 | 59.62 | 25.82 | 10.80 | 3.76 | 66.42 | 10.64 | 26.53 | 4.75 |
| | Train | 347 | 40.35 | 10.66 | 38.90 | 10.09 | 71.41 | 9.87 | 26.32 | 4.89 |
| BRCA | Test | 281 | 0.00 | 67.07 | 0.00 | 32.93 | 62.00 | 13.19 | 25.51 | 4.88 |
| | Train | 82 | 0.00 | 50.53 | 0.00 | 49.47 | 60.07 | 13.65 | 25.57 | 5.49 |
| LUCA | Test | 219 | 53.42 | 20.55 | 15.07 | 10.96 | 66.39 | 10.13 | 26.71 | 5.01 |
| | Train | 908 | 20.48 | 29.96 | 26.32 | 23.24 | 65.08 | 10.39 | 25.69 | 4.87 |
| | Stage 1 Train | 144 | 27.78 | 22.92 | 27.78 | 21.53 | 69.74 | 8.68 | 25.69 | 4.97 |
| | Stage 1+2 Train | 235 | 27.66 | 22.55 | 28.09 | 21.70 | 68.55 | 9.20 | 25.94 | 4.97 |
| | Stage 2 Train | 91 | 27.47 | 21.98 | 28.57 | 21.98 | 66.67 | 9.67 | 26.34 | 4.95 |
| | Stage 3 Train | 176 | 28.41 | 22.16 | 27.84 | 21.59 | 67.60 | 9.75 | 26.49 | 6.07 |
| | Stage 4 Train | 275 | 23.27 | 26.91 | 24.00 | 25.82 | 68.49 | 8.83 | 25.30 | 4.54 |
| PRCA | Test | 295 | 49.03 | 0.00 | 50.97 | 0.00 | 67.07 | 8.93 | 27.13 | 4.21 |
| | Train | 569 | 50.09 | 0.00 | 49.91 | 0.00 | 61.46 | 12.86 | 26.75 | 4.32 |

Table S1: Demographics across different diseases, stages, and train/test sets.

#### B Peak ratio indices

| Index | Peak Ratio |
|---|---|
| 1 | $I_{1635}/I_{1654}$ |
| 2 | $I_{1546}/I_{1655}$ |
| 3 | $I_{1655}/(I_{1655}+I_{1548})$ |
| 4 | $I_{1684}/(I_{1655}+I_{1548})$ |
| 5 | $I_{1515}/(I_{1655}+I_{1548})$ |
| 6 | $I_{2959}/I_{2931}$ |
| 7 | $(I_{2855}+I_{2927})/(I_{2962}+I_{2871})$ |
| 8 | $(I_{2851}+I_{2927})/(I_{1655}+I_{1548})$ |
| 9 | $I_{1239}/(I_{2851}+I_{2927})$ |
| 10 | $I_{1741}/I_{1640}$ |
| 11 | $I_{1740}/I_{1400}$ |
| 12 | $I_{2852}/I_{1400}$ |
| 13 | $I_{1450}/I_{1539}$ |
| 14 | $I_{1240}/I_{1517}$ |
| 15 | $I_{1045}/I_{1545}$ |
| 16 | $I_{1080}/I_{1550}$ |
| 17 | $I_{1060}/I_{1230}$ |
| 18 | $I_{1170}/I_{1080}$ |
| 19 | $I_{1030}/I_{1080}$ |
| 20 | $I_{1080}/I_{1243}$ |
| 21 | $I_{1587}/(I_{1655}+I_{1548})$ |
| 22 | $I_{1156}/I_{1171}$ |
| 23 | $I_{1243}/I_{1314}$ |
| 24 | $I_{1453}/I_{1400}$ |

Table S2: List of Peak Ratios and assigned indices in Figure 2

#### C AUCs for all FTIR datasets

| split | datatype | cohort | stage | testtype | metric | sex | auc | std auc |
|---|---|---|---|---|---|---|---|---|
| sex&disease | mvg | blca | - | cv | label | 0&1 | 77.36 | 6.48 |
| | | | | | sex | 0&1 | 93.79 | 4.74 |
| | | brca | - | cv | label | 0&1 | 86.22 | 6.55 |
| | | | | | sex | 0&1 | - | - |
| | | luca | - | cv | label | 0&1 | 93.09 | 2.19 |
| | | | | | sex | 0&1 | 94.17 | 2.15 |
| | | | 1 | cv | label | 0&1 | 87.6 | 9.23 |
| | | | | | sex | 0&1 | 92.77 | 6.5 |
| | | | 1&2 | cv | label | 0&1 | 85.09 | 9.02 |
| | | | | | sex | 0&1 | 94.5 | 3.75 |
| | | | 2 | cv | label | 0&1 | 86.94 | 10.54 |
| | | | | | sex | 0&1 | 88.25 | 12.8 |
| | | | 3 | cv | label | 0&1 | 91.08 | 7.13 |
| | | | | | sex | 0&1 | 92.44 | 5.16 |
| | | | 4 | cv | label | 0&1 | 94.41 | 4.09 |
| | | | | | sex | 0&1 | 92.44 | 4.52 |
| | | prca | - | cv | label | 0&1 | 83.21 | 5.78 |
| | | | | | sex | 0&1 | - | - |
| | | blca | - | test | label | 0&1 | 61.43 | - |
| | | | | | sex | 0&1 | 78.84 | - |
| | | brca | - | test | label | 0&1 | 50.17 | - |
| | | | | | sex | 0&1 | - | - |
| | | luca | - | test | label | 0&1 | 78.14 | - |
| | | | | | sex | 0&1 | 90.11 | - |
| | | prca | - | test | label | 0&1 | 60.27 | - |
| | | | | | sex | 0&1 | - | - |
| | real | blca | - | cv | label | 0&1 | 69.16 | 8.44 |
| | | | | | sex | 0&1 | 90.81 | 4.72 |
| | | brca | - | cv | label | 0&1 | 76.49 | 7.45 |
| | | | | | sex | 0&1 | - | - |
| | | luca | - | cv | label | 0&1 | 89.88 | 3.23 |
| | | | | | sex | 0&1 | 91.57 | 2.58 |
| | | | 1 | cv | label | 0&1 | 67.95 | 13.81 |
| | | | | | sex | 0&1 | 88.72 | 8.82 |
| | | | 1&2 | cv | label | 0&1 | 75.02 | 10.83 |
| | | | | | sex | 0&1 | 91.07 | 6.45 |
| | | | 2 | cv | label | 0&1 | 78.3 | 14.9 |
| | | | | | sex | 0&1 | 85.13 | 12.74 |
| | | | 3 | cv | label | 0&1 | 86.32 | 9.38 |
| | | | | | sex | 0&1 | 84.08 | 9.77 |
| | | | 4 | cv | label | 0&1 | 91.64 | 5.11 |
| | | | | | sex | 0&1 | 84.49 | 7.18 |
| | | prca | - | cv | label | 0&1 | 74.45 | 6.07 |
| | | | | | sex | 0&1 | - | - |
| | | blca | - | test | label | 0&1 | 61.5 | - |
| | | | | | sex | 0&1 | 89.13 | - |
| | | brca | - | test | label | 0&1 | 60.07 | - |
| | | | | | sex | 0&1 | - | - |
| | | luca | - | test | label | 0&1 | 85.91 | - |
| | | | | | sex | 0&1 | 92.68 | - |
| | | prca | - | test | label | 0&1 | 66.83 | - |
| | | | | | sex | 0&1 | - | - |
| | mvg | blca | - | cv | label | 0 | 80.21 | 7.18 |
| | | luca | - | cv | label | 0 | 93.94 | 3.73 |
| | | | 1 | cv | label | 0 | 92.29 | 8.99 |

Continued on next page

| split | datatype | cohort | stage | testtype | metric | sex | auc | std auc |
|---|---|---|---|---|---|---|---|---|
| | | | 1&2 | cv | label | 0 | 89.32 | 9.36 |
| | | | 2 | cv | label | 0 | 78.12 | 20.19 |
| | | | 3 | cv | label | 0 | 90.22 | 11.76 |
| | | | 4 | cv | label | 0 | 94.66 | 6.72 |
| | | prca | - | cv | label | 0 | 83.92 | 4.4 |
| | | blca | - | test | label | 0 | 63.06 | - |
| | | luca | - | test | label | 0 | 76.17 | - |
| | | prca | - | test | label | 0 | 60.27 | - |
| | real | blca | - | cv | label | 0 | 64.2 | 10.16 |
| | | luca | - | cv | label | 0 | 89.19 | 4.92 |
| | | | 1 | cv | label | 0 | 66.48 | 19.98 |
| | | | 1&2 | cv | label | 0 | 72.92 | 12.19 |
| | | | 2 | cv | label | 0 | 68.47 | 21.93 |
| | | | 3 | cv | label | 0 | 84.06 | 10.4 |
| | | | 4 | cv | label | 0 | 92.11 | 6.32 |
| | | prca | - | cv | label | 0 | 74.49 | 6.36 |
| | | blca | - | test | label | 0 | 62.38 | - |
| | | luca | - | test | label | 0 | 77.08 | - |
| | | prca | - | test | label | 0 | 66.83 | - |
| | mvg | blca | - | cv | label | 1 | 90.25 | 11.79 |
| | | brca | - | cv | label | 1 | 85.79 | 6.94 |
| | | luca | - | cv | label | 1 | 91.85 | 3.24 |
| | | | 1 | cv | label | 1 | 77.56 | 20.74 |
| | | | 1&2 | cv | label | 1 | 82.6 | 11.67 |
| | | | 2 | cv | label | 1 | 96.03 | 10.01 |
| | | | 3 | cv | label | 1 | 88.35 | 14.56 |
| | | | 4 | cv | label | 1 | 92.08 | 7.56 |
| | | blca | - | test | label | 1 | 47.05 | - |
| | | brca | - | test | label | 1 | 50.17 | - |
| | | luca | - | test | label | 1 | 93.7 | - |
| | real | blca | - | cv | label | 1 | 71.1 | 20.08 |
| | | brca | - | cv | label | 1 | 77.08 | 8.01 |
| | | luca | - | cv | label | 1 | 86.63 | 5.44 |
| | | | 1 | cv | label | 1 | 60.08 | 21.14 |
| | | | 1&2 | cv | label | 1 | 69.69 | 12.51 |
| | | | 2 | cv | label | 1 | 96.03 | 11.19 |
| | | | 3 | cv | label | 1 | 75.12 | 15.22 |
| | | | 4 | cv | label | 1 | 88.17 | 9.91 |
| | | blca | - | test | label | 1 | 49.09 | - |
| | | brca | - | test | label | 1 | 60.07 | - |
| | | luca | - | test | label | 1 | 90.83 | - |
| disease | mvg | blca | - | cv | label | _ | 78.73 | 7.19 |
| | | | | | sex | _ | - | - |
| | | brca | - | cv | label | _ | 87.17 | 6.58 |
| | | | | | sex | _ | - | - |
| | | luca | - | cv | label | _ | 89.61 | 2.88 |
| | | | | | sex | _ | - | - |
| | | | 1 | cv | label | _ | 84.86 | 9.54 |
| | | | | | sex | _ | - | - |
| | | | 1&2 | cv | label | _ | 85.95 | 6.34 |
| | | | | | sex | _ | - | - |
| | | | 2 | cv | label | _ | 85.85 | 12.31 |
| | | | | | sex | _ | - | - |
| | | | 3 | cv | label | _ | 93.45 | 5.5 |
| | | | | | sex | _ | - | - |
| | | | 4 | cv | label | _ | 95.89 | 3.07 |
| | | | | | sex | _ | - | - |
| | | prca | - | cv | label | _ | 83.91 | 5.28 |

Continued on next page

| split | datatype | cohort | stage | testtype | metric | sex | auc | std auc |
|---|---|---|---|---|---|---|---|---|
| | | | | | sex | _ | - | - |
| | | blca | - | test | label | _ | 60.14 | - |
| | | | | | sex | _ | - | - |
| | | brca | - | test | label | _ | 54.48 | - |
| | | | | | sex | _ | - | - |
| | | luca | - | test | label | _ | 81.61 | - |
| | | | | | sex | _ | - | - |
| | | prca | - | test | label | _ | 58.97 | - |
| | | | | | sex | _ | - | - |
| | real | blca | - | cv | label | _ | 69.16 | 8.44 |
| | | | | | sex | _ | 90.81 | 4.72 |
| | | brca | - | cv | label | _ | 76.49 | 7.45 |
| | | | | | sex | _ | - | - |
| | | luca | - | cv | label | _ | 89.88 | 3.23 |
| | | | | | sex | _ | 91.57 | 2.58 |
| | | | 1 | cv | label | _ | 67.95 | 13.81 |
| | | | | | sex | _ | 88.72 | 8.82 |
| | | | 1&2 | cv | label | _ | 75.02 | 10.83 |
| | | | | | sex | _ | 91.07 | 6.45 |
| | | | 2 | cv | label | _ | 78.3 | 14.9 |
| | | | | | sex | _ | 85.13 | 12.74 |
| | | | 3 | cv | label | _ | 86.32 | 9.38 |
| | | | | | sex | _ | 84.08 | 9.77 |
| | | | 4 | cv | label | _ | 91.64 | 5.11 |
| | | | | | sex | _ | 84.49 | 7.18 |
| | | prca | - | cv | label | _ | 74.45 | 6.07 |
| | | | | | sex | _ | - | - |
| | | blca | - | test | label | _ | 61.5 | - |
| | | | | | sex | _ | 89.13 | - |
| | | brca | - | test | label | _ | 60.07 | - |
| | | | | | sex | _ | - | - |
| | | luca | - | test | label | _ | 85.91 | - |
| | | | | | sex | _ | 92.68 | - |
| | | prca | - | test | label | _ | 66.83 | - |
| | | | | | sex | _ | - | - |

Table S3: AUCs for predicting the disease label and the sex on real and on simulated data on all cohorts.
